# Two centuries of change in the traits and origins of non-native vertebrates

**DOI:** 10.64898/2026.08.06.743192

**Authors:** Filipa Coutinho Soares, Joana Catarino, Matilde Mendes, Joana Ribeiro, Céline Bellard, Franz Essl, Chunlong Liu, Luís Reino, Hanno Seebens, César Capinha

## Abstract

Globalization is redistributing species worldwide, yet whether the traits and origins of non-native fauna have changed through time remains unclear. We combined global first-record data for non-native species from 1800–2019 with harmonized information on body size, diet, habitat use, native range characteristics, and climatic niche characteristics for 1,910 non-native mammals, birds, reptiles, amphibians, and freshwater fishes. Across most groups, species recorded earlier originated disproportionately from higher latitudes, occupied broader native ranges, and had wider thermal niches. More recent first records increasingly involve species from warmer, lower-latitude regions with smaller and more restricted native distributions. Body size also declined through time in several groups, whereas shifts in diet and habitat use were more taxon-specific. Temporal changes in introduction pathways partly explained these patterns: declines in deliberate release and production-related pathways, together with increases in pet and ornamental pathways, were associated with shifts toward smaller-bodied, lower-latitude, and more range-restricted species. These results indicate that the functional and biogeographic composition of non-native vertebrates has been progressively reshaped over the past two centuries, weakening the historical dominance of widespread temperate species and increasingly incorporating tropical and range-restricted fauna into global redistribution. Anticipating future biological invasions will therefore require attention not only to the number of species being transported, but also to how the characteristics and pathways of transported species are changing through time.

## Introduction

The expansion of transport networks and the growing movement of people and goods are driving an unprecedented redistribution of species beyond their native ranges (IPBES, 2023). Many introduced species establish self-sustaining populations, becoming non-native species in recipient regions and eventually altering ecological communities, ecosystem functioning, economic activities, and human well-being (Bellard et al., 2021; Diagne et al., 2021). The global accumulation of non-native species has continued to accelerate over the past two centuries, particularly in recent decades, despite increasing scientific attention and strengthened biosecurity (Seebens et al., 2025). Anticipating the ecological consequences of this ongoing redistribution requires understanding not only how many species are transported, but also whether the characteristics of the species entering the non-native pool are changing through time.

Species traits influence multiple stages of the introduction and invasion process. They may affect a species’ likelihood of being selected for deliberate introduction or accidentally transported, as well as its ability to survive transport and release, establish self-sustaining populations, spread, and cause ecological impacts (Capellini et al., 2015; Hodgins et al., 2018; Redding et al., 2019; Enders et al., 2020; Evans et al., 2021). Trait-based approaches have therefore become central to invasion ecology. Most studies, however, have focused on identifying characteristics that distinguish non-native from native species or successful invaders from species that fail to establish or spread (e.g., Allen et al., 2017; Mathakutha et al., 2019). These comparisons generally treat the pool of non-native species as temporally invariant (but see Dyer et al., 2017 for birds). Consequently, it remains unclear whether the characteristics of animals redistributed outside their native ranges by humans today differ systematically from those introduced during earlier phases of global environmental and socioeconomic change, and whether such changes are consistent across major vertebrate groups. If the composition of the non-native species pool is changing, then risk assessments based primarily on historical introductions may become less effective at anticipating future invaders.

The species transported by humans are unlikely to represent a random sample of global biodiversity. Human preferences, commercial uses, transport opportunities, and the ecological requirements of species can all influence which animals are moved beyond their native ranges (Long, 2003; Hulme et al., 2008; Su et al., 2016; Blackburn et al., 2017; Reino et al., 2017; Hinz et al., 2019; Bernery et al., 2024; Liu et al., 2025). These influences have themselves changed substantially through colonial expansion, industrialization, and modern globalization. Nineteenth-century acclimatization movements, for example, promoted the deliberate introduction of familiar European birds and mammals into settler regions, while fisheries programmes distributed a small set of commercially valued fishes (Duncan et al., 2003; Casal, 2006; McLeod & Saunders, 2014; Blackburn et al., 2015). More recently, the expansion of international trade, transport networks, and consumer markets has increased the geographic reach and taxonomic diversity of transported animals, including reptiles, amphibians, fishes, birds, and mammals associated with the pet, aquarium, and wildlife trades (Kraus, 2009; Hughes et al., 2023). These historical changes in introduction pathways provide reason to expect corresponding shifts in the functional and biogeographic characteristics of newly recorded non-native fauna. Yet, it remains unknown whether the traits of non-native vertebrates have changed systematically through time, and whether such changes are associated with shifting introduction pathways, at a global scale and across major vertebrate groups.

Here, we quantify temporal changes in the traits of non-native terrestrial and freshwater vertebrates worldwide. Using the Alien Species First Records Database (Seebens et al., 2023), we identified the earliest recorded introduction year of each non-native species at the global scale and combined these records with harmonized functional and biogeographic trait information for more than 1,900 mammals, birds, reptiles, amphibians, and freshwater fishes. This comparative framework allowed us to evaluate whether long-term changes in the characteristics of non-native species are consistent across major vertebrate groups. The traits considered describe body size, diet, habitat use, native range extent and geographic position, human population density within the native range, and native climatic conditions and niche breadths. We first tested whether these characteristics showed directional trends among species first recorded between 1800 and 2019. Then, we assessed whether species traits predicted introduction timing across taxonomic groups after accounting for geographic differences among recipient regions and examined whether these relationships were nonlinear. Finally, we evaluated whether changes in broad introduction pathway classes were associated with the main temporal shifts. By reconstructing two centuries of change in the characteristics and pathways of non-native vertebrates, we provide a dynamic perspective on how both traits and pathways of non-native vertebrates have changed through successive phases of globalization, with implications for anticipating future biological invasions.

## Results

### Temporal trends in traits of non-native vertebrates

We analyzed 1,910 non-native vertebrate species first recorded globally between 1800 and 2019, including 234 mammals, 931 birds, 198 reptiles, 100 amphibians, and 445 freshwater fishes. Each species was represented by its earliest known global record. We assessed temporal trends in species traits using Kendall’s rank correlations between raw species-level trait values and the first-record year.

The traits of newly recorded non-native species changed markedly through time (Fig. 1; SI Appendix, Fig. S1 and Tables S2–S6). The most consistent cross-taxon changes concerned absolute latitude of species’ native ranges, and species native range area and number of native continents. Except for amphibians, the absolute latitude of species’ native ranges declined over time, while native range area and the number of native origin continents declined in mammals, birds, and freshwater fishes. Reptiles also showed a decline in human population density within their native ranges, particularly from the mid-eighteenth to the mid-nineteenth century (SI Appendix, Fig. S1). Human population density showed no consistent temporal trend in the other groups.

**Figure 1.**
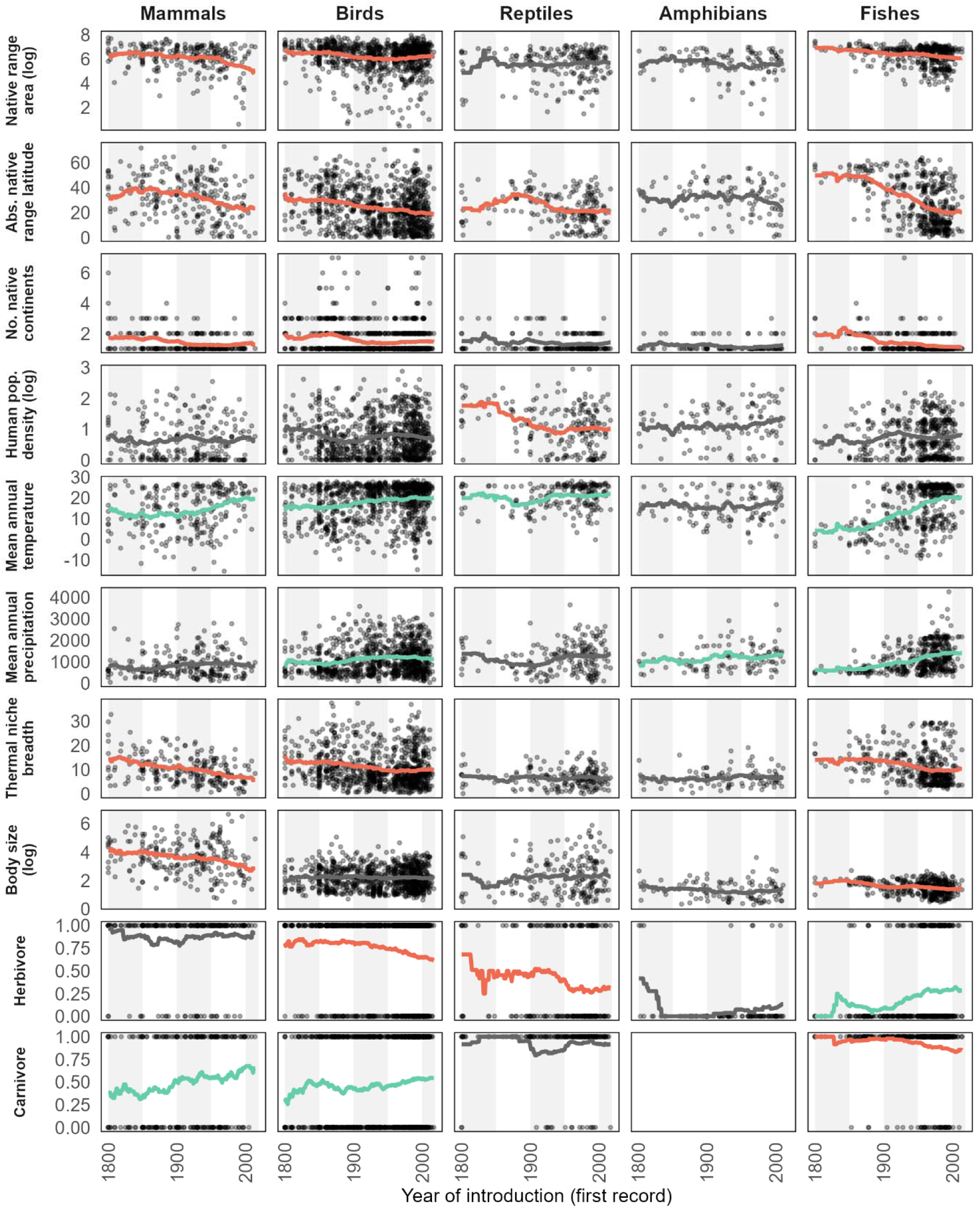
Species-level trait values for the most consistent cross-taxon changes, plotted against the first-recorded year for each taxonomic group. Lines represent smoothed temporal trends based on a centered 50-year moving average of annual mean trait values, highlighting long-term patterns while reducing short-term variation. For binary traits, trends represent temporal changes in the proportion of newly recorded non-native species exhibiting the respective trait. Shaded vertical bands indicate 50-year periods used to visualize temporal structure. Points represent individual species according to their earliest known global first record. Line colors indicate the direction and significance of Kendall’s rank correlations: green denotes significant positive relationships, red denotes significant negative relationships, and black the non-significant relationships (statistical significance defined at *P* value<0.05). Some traits were log-transformed for visualization.

Climatic niche characteristics changed alongside geographic origins (Fig. 1; SI Appendix, Fig. S1). Mean annual temperature within native ranges increased over time in mammals, birds, reptiles, and freshwater fishes. The increase was especially pronounced in fishes from the late eighteenth century onward, whereas mammals and birds showed more gradual warming through the nineteenth century, and reptiles increased mainly after a mid-eighteenth-century decline. Thermal niche breadth declined in mammals, birds, and fishes, with the clearest reductions occurring progressively through the nineteenth century. Mean annual precipitation and precipitation niche breadth increased in birds, amphibians, and fishes, generally becoming more apparent from the mid-nineteenth century onward, although these patterns were less pronounced than those for temperature.

Functional traits showed more taxon-specific changes (Fig. 1; SI Appendix, Fig. S1). Body size declined in mammals and freshwater fishes, gradually through the nineteenth and twentieth centuries in mammals and mainly between the late nineteenth and mid-twentieth centuries in fishes. Herbivores declined among birds and reptiles but increased among fish. The decline was gradual and strongest after approximately 1950 in birds, whereas reptiles showed a pronounced reduction during the nineteenth century. In fish, herbivores became increasingly represented from the early nineteenth century onward, particularly after mid-century. Carnivores increased among mammals and birds but declined among fish. Omnivores increased only among mammals, mainly from the early to late nineteenth century. Habitat associations were otherwise largely stable, except for an increasing proportion of freshwater-associated birds toward the end of the nineteenth century.

### Species traits predict introduction timing

To evaluate whether species traits jointly explained variation in introduction timing, we fitted separate linear mixed-effects models for each taxonomic group, estimating first-record year as a function of species traits while accounting for variation among recipient continents. We complemented these analyses with boosted regression trees (BRTs) to quantify relative predictor importance and identify nonlinear relationships. Both approaches consistently identified native range geography (absolute native range latitude and native range area) and climatic characteristics (mean annual temperature and precipitation, thermal niche breadth) as the strongest correlates of introduction timing across taxa (Fig. 2; SI Appendix, Fig. S2–S6). These results indicate a long-term shift from widespread, higher-latitude, and broad-niched vertebrates toward lower-latitude, range-restricted, and narrower-niched species, whereas changes in functional traits were more taxon-specific.

**Figure 2.**
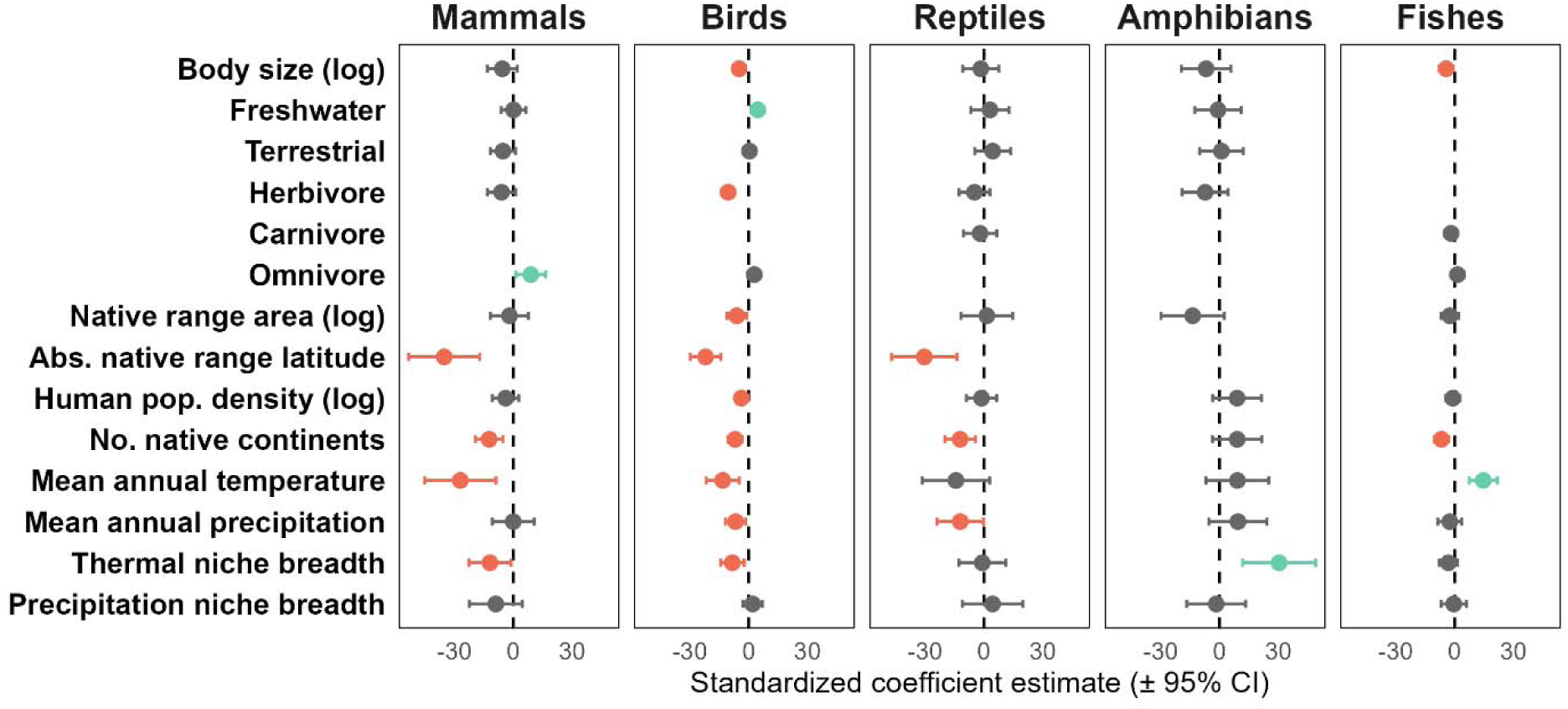
Associations between species traits and year of first recorded introduction across taxonomic groups. Points represent standardized fixed-effect coefficients from linear mixed-effects models, and horizontal bars indicate 95% confidence intervals. Colors denote the direction and statistical significance of associations (*P* value<0.05): green indicates significant positive associations (e.g., higher values associated with recent introductions), red indicates significant negative associations (e.g., higher values associated with earlier introductions), and grey indicates non-significant associations.

Absolute native range latitude and the number of native continents were the most recurrent predictors in mixed-effects models: species from higher latitudes were recorded earlier in mammals, birds, and reptiles, while species occupying more native continents were associated with earlier records in mammals, birds, reptiles, and fishes (Fig. 2). In contrast, native range area showed a weaker and more taxon-specific association, with larger-ranged species being recorded earlier only in birds. Climatic niche characteristics were also important but more taxon-dependent: broader thermal niches were associated with earlier records in mammals and birds but later records in amphibians, whereas warmer native climates were associated with earlier records in mammals and birds but later records in fishes. Functional traits showed fewer consistent relationships in mixed-effects models, although larger body size was linked to earlier records in birds and fishes, freshwater-associated birds and omnivorous mammals were recorded later, and herbivorous birds were recorded earlier.

Relative importance values from the BRTs broadly supported the mixed-effects models by emphasizing native range, climatic, and body-size traits, although the most influential predictors differed among taxa (SI Appendix, Tables S13–S17). Thermal niche breadth was the strongest predictor of introduction timing in mammals and fishes and was also important in birds and amphibians, whereas native range latitude was most influential in birds and contributed substantially to mammals and reptiles. BRTs further highlighted human population density in reptiles, mammals, and birds, body size in mammals, reptiles, and fishes, herbivory in birds, mean annual temperature in fishes, and mean annual precipitation in amphibians. Partial-dependence plots (SI Appendix, Fig. S2–S6) further showed that these relationships varied through time among vertebrate groups. In mammals and birds, broader thermal niches, higher native range latitudes, and greater human population densities were associated with earlier records, with the strongest temporal shifts occurring earlier in mammals, from the early 1900s to around 1930, and slightly later in birds, from the late 1920s to early 1940s. Larger body size was also associated with earlier records in mammals, while herbivory was associated with earlier records in birds. Reptiles showed later and more abrupt responses, mainly from the late 1930s to around 1950, with higher latitude, higher human population density, and more native continents generally associated with earlier records. In fishes, broader thermal niches, larger body size, and broader native ranges were associated with earlier records, whereas warmer native climates were associated with later records, showing strongest temporal shifts from the mid-1940s to mid-1950s. Amphibians showed little change in region-adjusted first-record year.

The two modelling approaches also showed comparable performance across taxa. Marginal R² values from the mixed-effects models ranged from 0.13 for birds and reptiles to 0.28 in mammals, increasing to conditional R² values between 0.23 and 0.36, after accounting for recipient continent (SI Appendix, Tables S7–S12). The larger difference between marginal and conditional R² values for birds and reptiles indicates a stronger contribution of recipient continent to explaining first-record timing. Similarly, BRT predictive performance was higher for birds (relative absolute error = 0.80) and fishes (0.83), intermediate for mammals (0.90) and reptiles (0.91), and lowest for amphibians (0.97), indicating limited predictive power for the latter.

### Shifts in introduction pathways partly account for temporal trait change

We finally assessed whether temporal changes in introduction pathways among first-recorded non-native species contributed to the trait shifts identified above. We focused on four key traits that showed the most consistent temporal patterns in the previous analyses: body size, absolute native range latitude, number of native continents, and thermal niche breadth. Pathway subcategories were grouped into six non-mutually exclusive classes: deliberate release, production escape, pet and ornamental keeping, biological contaminants, transport stowaways, and transport corridors.

Pathway composition of first-recorded species differed markedly among taxa, while temporal changes were generally gradual (Fig. 3; SI Appendix, Fig. S7). First-recorded species associated with deliberate release declined after 1900 and 1950 in birds, fish, and mammals, whereas those associated with pet and ornamental pathways increased or maintained in all taxa (SI Appendix, Fig. S7). Fish introductions associated with production escape remained stable or increased slightly before declining after 2000, while introductions via transport corridors remained stable before also declining. Reptiles and amphibians were increasingly dominated by pet and ornamental pathways (SI Appendix, Fig. S7).

**Figure 3.**
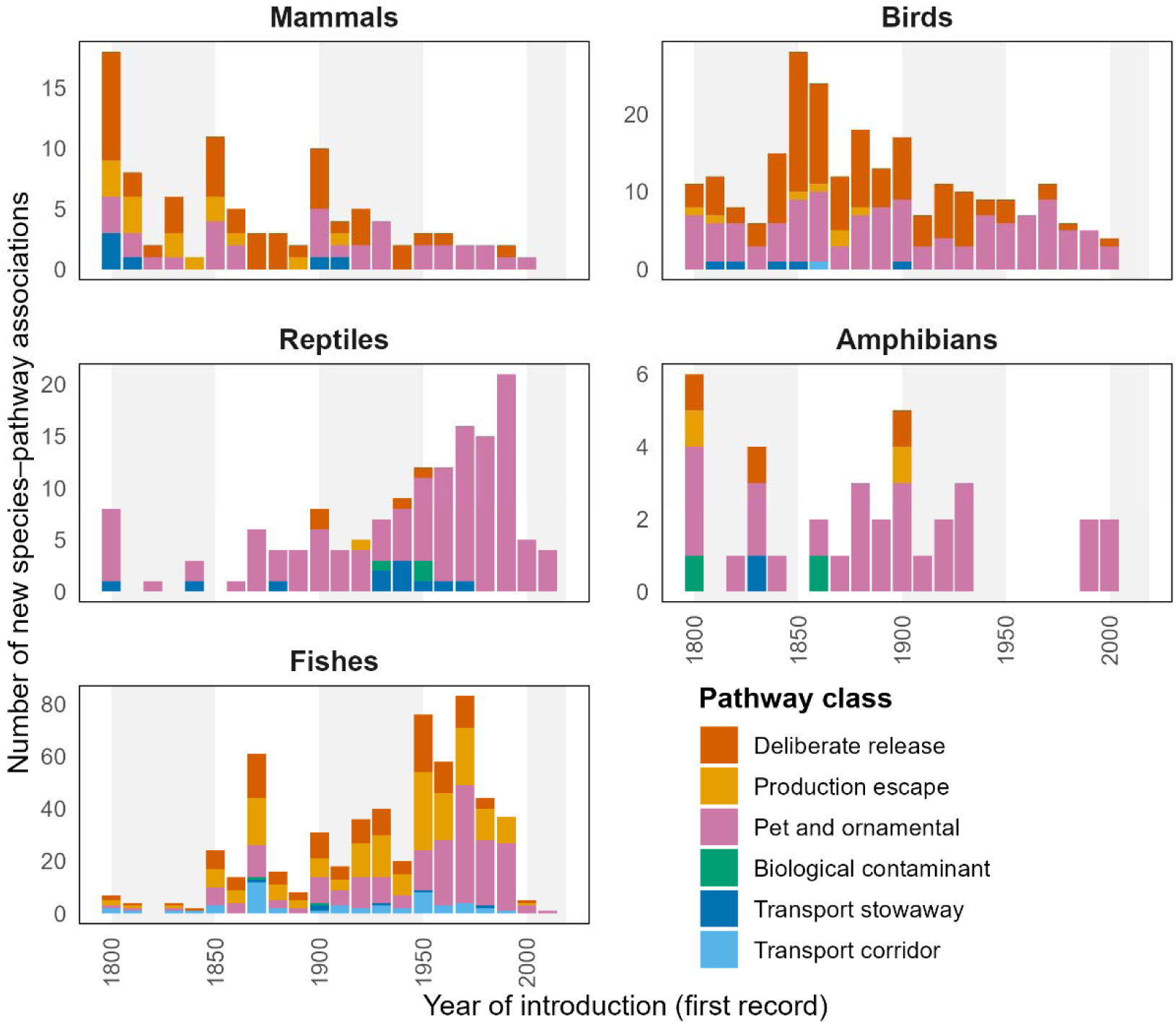
Number of new species–pathway associations per decade for each taxonomic group, based on the earliest global first record of each species between 1800 and 2019. Colours indicate six broad introduction pathway classes: deliberate release, biological contaminant, pet and ornamental, transport corridor, production escape, and transport stowaway. Because pathway classes are non-mutually exclusive, individual species may contribute to more than one pathway category (786 species: 28 amphibians, 210 birds, 344 freshwater fishes, 71 mammals, and 133 reptiles).

Mediation analyses indicated that these pathway shifts partly accounted for changes in body size and native range latitude, but less consistently for native distribution breadth (absolute latitude and number of native continents) or thermal niche breadth (Fig. 4). In birds, declines in deliberate release and increases in pet and ornamental pathways were associated with shifts toward smaller-bodied and lower-latitude species. In fishes, declines in deliberate release, production escape, and transport corridors were associated with lower-latitude species, with production escape also linked to narrower thermal niches. In mammals, declines in deliberate release and production escape were associated mainly with smaller body size and, more weakly, lower latitude or narrower thermal breadth. Reptiles showed little evidence of pathway-mediated trait change, and amphibian tests were limited by sample size.

**Figure 4.**
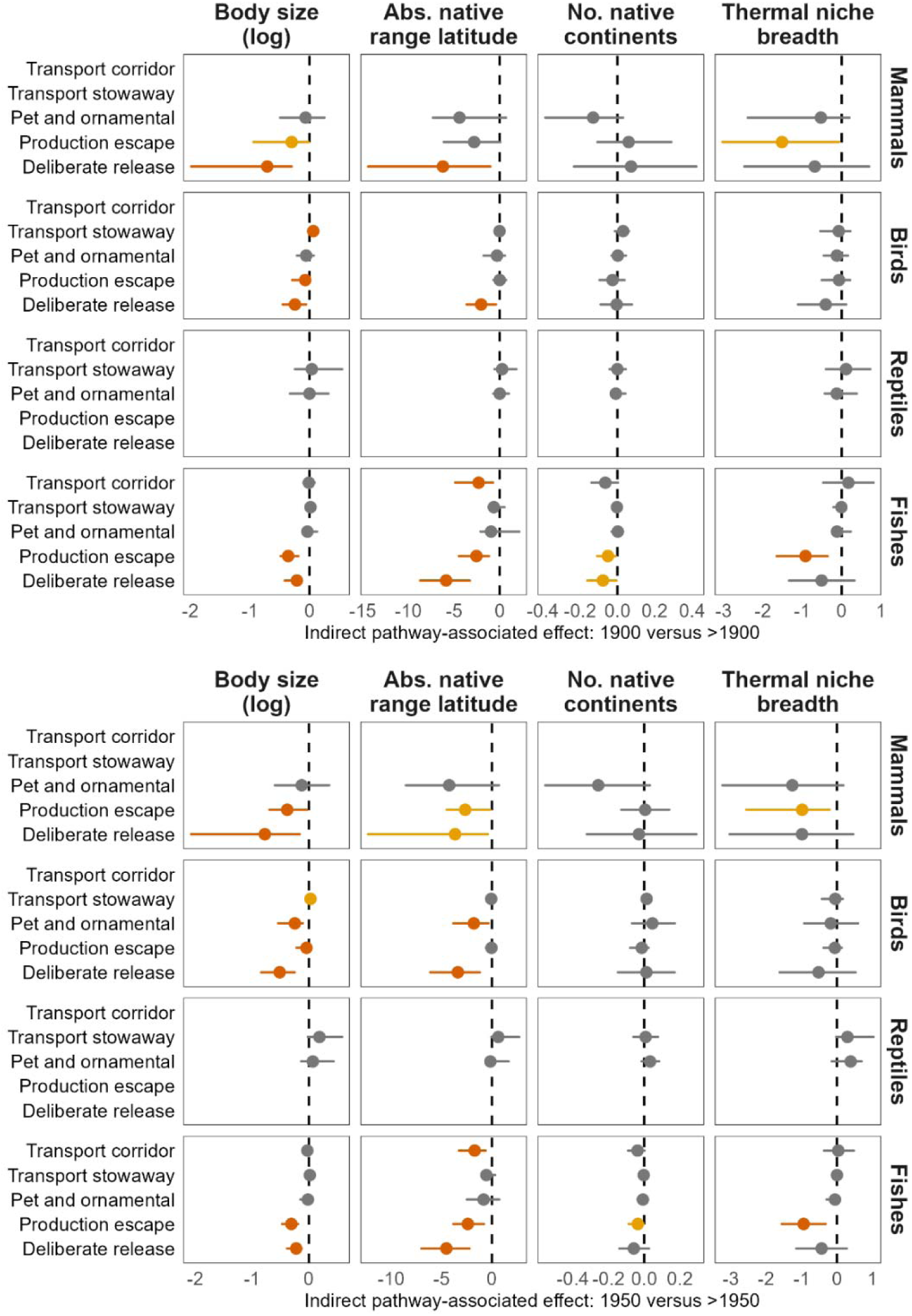
Pathway-associated mediation effects on key trait shifts across historical cut-offs of 1900 (upper panel) and 1950 (lower panel). Points show average indirect effects for each pathway, taxonomic group, and trait, whereas horizontal lines indicate bootstrap 95% confidence intervals. Vertical dashed lines indicate zero effect. Negative values indicate that pathway turnover was associated with lower trait values after the cut-off, whereas positive values indicate association with higher trait values. Colours indicate statistical support after Benjamini–Hochberg correction, nominal significance, or non-significance.

## Discussion

Our results reveal consistent temporal shifts in the traits of non-native vertebrates first recorded worldwide over the past two centuries. Earlier introductions were associated with species from higher latitudes, with broad native distributions and wider thermal niche breadths, whereas more recent records increasingly involved species originating from lower latitudes with smaller native ranges and narrower thermal niches. Several vertebrate groups also showed a trend toward smaller-bodied species through time.

The clearest temporal change involved the geographic origins of non-native vertebrates. Nineteenth- and early twentieth-century introductions were strongly shaped by European colonial expansion, acclimatization movements, game releases, and fisheries development, which repeatedly transported familiar temperate species to overseas territories (Duncan et al., 2003; Clout & Russell, 2008; Blackburn et al., 2015; Dyer et al., 2017). These activities favored a relatively restricted pool of widespread, robust, and often economically or culturally valued species. In contrast, modern transport networks and global consumer markets connect a much larger set of source regions (Seebens et al., 2018) and facilitate the movement of reptiles, amphibians, ornamental fishes, birds, and mammals through the pet, aquarium, and wildlife trades (Kraus, 2009; Lockwood et al., 2019; Li et al., 2023). Our finding that recent first records increasingly involve lower-latitude and geographically restricted species is consistent with this historical broadening of the geographic sources contributing to global species introductions.

Analyses of introduction pathways help explain these shifts. Declines in deliberate release and production-related pathways were consistently associated with shifts away from larger-bodied, higher-latitude, and broader-niched species, particularly in birds, fishes, and mammals. This agrees with the historical role of acclimatization societies, game releases, fisheries enhancement, and livestock or production systems in repeatedly moving familiar, often temperate species selected for their utility, availability, and perceived likelihood of establishment (Duncan et al., 2003; Casal, 2006; Clout & Russell, 2008; Blackburn et al., 2015). Conversely, the increasing representation of pet and ornamental pathways, especially in birds, reptiles, amphibians, and mammals, reflects the growing importance of consumer-driven wildlife trade in contemporary introductions. Such pathways can favor species valued for novelty, appearance, rarity, or ease of keeping (Jarić et al., 2020) rather than for direct utilitarian purposes and are known to draw heavily on tropical and subtropical faunas (Kraus, 2009; Lockwood et al., 2019; Altherr & Lameter, 2020; Toomes et al., 2020; Li et al., 2023). They may also favor smaller-bodied species that are easier to transport and maintain in captivity, consistent with the temporal declines in body size observed in mammals, birds, and fishes (e.g., Long, 2003; Dyer et al., 2017). Likewise, the declining representation of herbivore birds among more recent first records may reflect the transition away from historical introductions for food production and game toward species traded primarily as pets or ornamentals. In our analyses, the rise of pet and ornamental pathways was most clearly associated with lower-latitude origins in birds, while reptiles and amphibians were already strongly dominated by this pathway. Together, these results suggest that changes in introduction pathways have contributed to the long-term shift from introductions dominated by widespread temperate species toward more geographically diverse, market-driven movements of tropical and often more range-restricted vertebrates.

The nonlinear analyses further suggest that these transitions did not occur simultaneously across vertebrate groups. Trait shifts emerged earlier in mammals and birds and later in reptiles and freshwater fishes, broadly matching the historical sequence in which different taxa became prominent in global trade (Seebens et al., 2025). Mammals and birds featured heavily in colonial, agricultural, hunting, and acclimatization programs during the nineteenth and early twentieth centuries, whereas reptiles, ornamental fishes, and other taxa associated with captivity and trade became increasingly important only with the later expansion of international pet and aquarium markets around the middle of the twentieth century. Although these periods should not be interpreted as formal change points, they suggest that successive waves of globalization reshaped the traits of introduced vertebrates at different times across taxonomic groups.

The increasing representation of range-restricted species is especially noteworthy. Historically widespread species may have been more likely to encounter transport networks and to be repeatedly moved for food, hunting, fisheries, or other utilitarian purposes (e.g., Long, 2003; Blackburn et al., 2017). Contemporary markets, by contrast, can favor novelty, rarity, and taxonomic distinctiveness, particularly in the exotic pet and wildlife trades (Courchamp et al., 2006; Altherr & Lameter, 2020; Toomes et al., 2020). This may expose species with small native distributions to two simultaneous pressures: harvesting and trade within their native ranges, and introduction beyond them. The redistribution of range-restricted species is therefore relevant not only to invasion risk but also to conservation in source regions (Tedeschi et al., 2025).

The decline in thermal niche breadth suggests that contemporary introductions increasingly include species with more restricted climatic tolerances. Earlier long-distance transport and release may have imposed strong filters favoring generalists capable of surviving slow transport, handling, and establishment in unfamiliar environments (Blackburn et al., 2017; Dyer et al., 2017). Faster transport, improved husbandry, captive breeding, and specialized trade networks may allow species with a broader range of ecological strategies to enter global transport pathways. However, the patterns observed in first records are likely to reflect multiple filters operating across the introduction process, including transport, establishment, and detection. Recipient environments may therefore also contribute to determining which introduced species are ultimately recorded, although disentangling these mechanisms was beyond the scope of this study (Enders et al., 2020). Narrower thermal niches may also reduce establishment probability in many recipient regions, potentially resulting in a more heterogeneous pool of introductions in which many species may fail to establish, while those encountering suitable environmental conditions introduce traits and ecological interactions that were less common among historical invaders.

Our findings have implications for invasion forecasting and risk assessment. Many vertebrate species risk profiles incorporate historical invasion success, particularly establishment elsewhere, propagule/colonization pressure, and climate matching. As a result, they often prioritize species that have already demonstrated broad geographic, ecological, or climatic tolerances, or repeated success under human-mediated introduction pathways (e.g., Bomford et al., 2010; Dyer et al., 2016; Biancolini & Rondinini, 2025). Yet the species now entering transport networks increasingly depart from this historical profile. Approaches trained primarily on past invaders may consequently become less transferable to future introductions if the composition of transported species continues to shift. Horizon scanning and biosecurity assessment should therefore account not only for changes in introduction rates, but also for shifts in source regions, trade preferences, pathway composition, and the functional and biogeographic characteristics of transported species.

Several limitations qualify these interpretations. First-record years represent the earliest known detections rather than actual introduction dates, and detection lags vary among taxa, regions, and historical periods (Crooks et al., 2005; Zenetos et al., 2019). The analyses also combine established and non-established species and therefore characterize the transported and recorded species pool rather than establishment success. Pathway information was available at broad, non-mutually exclusive classes, meaning that pathway effects should be interpreted as pathway-associated contributions rather than exclusive causal mechanisms (Saul et al., 2017). In addition, mediation analyses were observational and should not be read as definitive evidence of causation.

Despite these limitations, the consistency of results among analytical approaches reveals a major change in the human-mediated redistribution of vertebrates. Globalization has progressively shifted introductions away from a historical concentration on widespread, temperate species broad niches and toward fauna originating from lower latitudes and possessing more restricted distributions and thermal niches. This shift reflects both the decline of historically important utilitarian pathways and the rise of more diverse, market-driven forms of wildlife movement. Anticipating future invasions will therefore require attention not only to how many species are transported, but also to how the identity, pathways, and characteristics of those species are changing.

## Materials and Methods

### First record data

We used the Alien Species First Records Database v3.1 (Seebens et al., 2023), which records the earliest known year in which non-native species were detected in the wild across 296 geographic regions. The database includes more than 75,000 first-record entries for over 24,000 non-native species and covers records at national and subnational levels. Because we aimed to quantify long-term changes in the traits of non-native vertebrates, rather than establishment success or impact, we retained records irrespective of invasion status, including established, non-established, eradicated, and extinct non-native species.

We restricted the analysis to terrestrial and freshwater vertebrates: mammals, birds, reptiles, amphibians, and freshwater fishes. Marine species were excluded using habitat classifications from FishBase, based on freshwater, brackish, and saltwater occurrence fields (Froese & Pauly, 2025). A small number of marine mammals and seabirds were also identified manually and removed from the dataset. The final dataset comprised 1,971 non-native vertebrate species.

### Species traits

We compiled data on 14 functional and biogeographical species traits, describing body size, diet (herbivore, carnivore, omnivore), habitat use (terrestrial, freshwater), native range characteristics (native range area, absolute native range latitude, number of native continents, human population density), and native climatic niche characteristics (mean annual temperature and precipitation, thermal and precipitation niche breadths; SI Appendix, Table S1). Trait information was obtained from taxon-specific trait databases: BirdLife International (Birdlife, 2025) and AVONET (Tobias et al., 2022) for birds; AmphiBIO (Oliveira et al., 2017) for amphibians; PHYLACINE (Faurby et al., 2018) for mammals; ReptTraits (v1.2; Oskyrko et al., 2024) for reptiles; and FishBase (Froese & Pauly, 2025) together with ITOFF (Invasive Traits of Freshwater Fish; Jessop et al., 2023) for freshwater fishes. Where trait values were missing, we supplemented the dataset with targeted literature searches when available.

Native range traits were derived from spatial range data for amphibians and mammals from the IUCN Red List (IUCN, 2025), birds from BirdLife (BirdLife, 2025), reptiles from the Global Assessment of Reptile Distributions (Roll et al., 2017), and freshwater fishes from Tedesco et al. (2017). We retained only native or reintroduced ranges, excluding non-native, vagrant, uncertain, and assisted-colonization areas. For each species, we calculated native range area, mean absolute native range latitude, and number of native continents. Native ranges were also overlaid with WorldClim v2.1 climate data (Fick & Hijmans, 2017) to estimate thermal niche breadth and summarize multivariate climatic conditions, and with HYDE human population density for 1975 (Goldewijk et al., 2011) to estimate median human population density within the native range. Spatial analyses were conducted in R (R Core Team, 2025), using equal-area projections for area calculations and supplementary sources for a small number of missing mammal ranges or human population density values.

Species lacking data for one or more trait variables were excluded from analyses. Complete trait information was available for 1,931 species, representing about 98% of all non-marine vertebrates included in the Alien Species First Record Database.

### Introduction Pathways

Introduction pathways were assigned using the harmonized dataset of Saul et al. (2017), which, to our knowledge, provides the most comprehensive standardized source of alien-species pathway information. We used the Convention on Biological Diversity pathway subcategories (CBD, 2014) as the starting classification and matched species names in this dataset to those in our vertebrate first-record dataset.

To reduce sparsity and align pathway information with hypothesized selective filters on species traits, we grouped CBD subcategories into six broad, non-mutually exclusive pathway classes: deliberate release, production escape, pet and ornamental keeping, biological contaminant, transport stowaway, and transport corridor (Hulme et al., 2008; Saul et al., 2017). These classes distinguish pathways expected to differ in trait selectivity, from utilitarian releases and production systems to consumer-driven pet and ornamental trade or accidental transport (Kraus, 2009; Blackburn et al., 2017; Lockwood et al., 2019). Each species was coded as present or absent in each pathway class and then linked to its earliest global first record from 1800 onward. The final pathway dataset included 786 species: 28 amphibians, 210 birds, 344 freshwater fishes, 71 mammals, and 133 reptiles.

### Temporal trends of non-native species traits

To assess whether functional and biogeographic traits exhibited directional temporal trends in the global pool of non-native vertebrates, we quantified the relationship between each trait and the year of the first recorded introduction using Kendall’s rank correlation coefficient (*stats* package; R Core Team, 2025). Analyses were restricted to the period 1800-2019 and were based on the earliest global first-record year per species, ensuring that each species contributed only once. This approach captures temporal changes in the traits of first-record non-native species rather than changes driven by repeated introductions.

To visualize long-term dynamics, we constructed time series of trait values using annual means and applied a centered 50-year moving average. This smoothing was used exclusively for visualization, as Kendall’s rank correlation coefficient was calculated with raw data. For binary traits, annual trait values represent the proportion of species exhibiting the trait.

### Multivariate modelling of trait-time relationships

To test whether temporal trends in functional and biogeographic traits were consistent across taxonomic groups and geographic regions, we applied two complementary modelling approaches: linear mixed-effects models (LMMs) and boosted regression trees (BRTs) incorporating mixed-effects structure. All analyses were performed separately for each vertebrate taxonomic group (mammals, birds, reptiles, amphibians, and freshwater fishes) in R (v. 4.4.3; R Core Team, 2025).

#### Linear mixed-effects models

For each taxonomic group, we fitted a linear mixed-effects model with the year of first recorded introduction as the response variable, using the *lme4* package (Bates et al., 2015). Models included species functional and biogeographic traits as fixed effects (predictors) and geographic region (Asia, Europe, North America, Africa, Oceania, Central and South America, Australia and New Zealand) as a random intercept, thereby accounting for non-independence among introduction events within regions and allowing baseline introduction timing to vary geographically.

Prior to modelling, continuous traits with strongly skewed distributions were log-transformed, notably body size, native range area, mean annual precipitation, thermal niche breadth, and precipitation niche breadth. Then, all continuous traits were standardized (mean = 0, SD = 1) to facilitate comparison of effect sizes, while binary traits were encoded as 0/1 variables. To account for multicollinearity among predictors, we applied the variance inflation factor (VIF; Fox et al., 2016) filtering using the *car* package, iteratively removing all predictors with VIF values greater than 10. This led to the removal of a few predictors from final models, namely diet-related categories (carnivores, herbivores, and omnivores).

Final models were fitted using restricted maximum likelihood (REML). Model performance was evaluated using marginal and conditional R² values via the *MuMIn* package (Barton & Barton, 2015), which represent, respectively, the variance explained by fixed effects alone and by both fixed and random effects combined (Nakagawa et al., 2017).

#### Boosted regression trees with mixed-effects structure

To capture potential non-linear relationships between introduction timing and species traits, we implemented boosted regression trees (BRT; Elith et al., 2008) using the *gbm* package (Ridgeway & Developers, 2026), while explicitly accounting for geographic structure.

Because standard BRTs do not accommodate random effects, we adopted a two-step approach. First, we fitted an intercept-only linear mixed-effects model with first-record year as the response variable and geographic region as a random intercept. This model estimated region-specific baseline differences in first-record timing (best linear unbiased predictions; Robinson, 1991), independent of species traits. These estimates were then subtracted from the observed first recorded introduction years to generate an adjusted response variable representing variation attributable to species traits after accounting for regional effects.

BRT models were subsequently fitted to this adjusted response using Gaussian error distributions and unstandardized trait predictors. Model parameters were set to 5,000 trees, an interaction depth of 4, a learning rate of 0.005, and a bag fraction of 0.8. Model performance was assessed using 10-fold cross-validation. Predictions were reconstructed by adding back the regional random effects, and predictive accuracy was quantified using the relative absolute error (RAE; Witten et al., 2005), which compares absolute prediction error with that of a baseline model predicting the mean observed first-record year. Lower values (below 1) indicate better predictive performance, whereas values approaching 1 indicate little improvement over simply predicting the average first-record year.

Final models were trained on the full dataset to estimate variable importance, expressed as relative influence scores. Partial dependence plots were generated for predictors with high relative influence (≥10%) to visualize their marginal effects on introduction timing while averaging over the influence of other predictors.

### Assessing Pathway Contributions to Trait Shifts

To examine temporal changes in species–pathway associations and to test whether pathway turnover was associated with shifts in key traits, we compared pathway composition before and after two historical cut-offs supported by Kendal- and BRT-based assessments, 1900 and 1950. Species first recorded up to and including each cut-off were contrasted with species first recorded after that cut-off. For each taxonomic group and pathway class, we estimated the relative frequency of pathway-associated species in each period. Because pathway classes were non-mutually exclusive, each pathway was analyzed separately.

We then used mediation analysis (Imai et al., 2010) to evaluate whether changes in pathway composition were associated with temporal shifts in key traits. Mediation analysis partitions the association between an exposure and an outcome into an indirect component acting through an intermediate variable and a remaining direct component (Imai et al., 2010; Tingley et al., 2014). Here, the period was the exposure, pathway presence was the mediator, and each trait was the outcome. For each taxonomic group, pathway, trait, and cut-off, using the *mediation* package (Tingley et al., 2014), we fitted a binomial mediator model testing whether pathway presence differed between periods and a linear outcome model testing whether trait values differed between periods after accounting for pathway presence. Indirect pathway-associated effects were estimated using nonparametric bootstrap resampling with 5,000 simulations. Analyses were omitted when the sample size or variation in period, pathway presence, or trait values was insufficient. *P* values were adjusted using the Benjamini–Hochberg procedure within each trait and cut-off (Tingley et al., 2014). Because the data are observational and pathways are broad and non-mutually exclusive, these analyses were interpreted as evidence of pathway-associated contributions to trait shifts, rather than definitive causal mediation.

## Acknowledgments

The authors acknowledge funding from *Fundação para a Ciência e a Tecnologia* (FCT) through the InvaSTOP project grant (https://doi.org/10.54499/2023.12533.PEX) and through funds to CEG/IGOT Research Unit (UID/00295/2025; https://doi.org/10.54499/UID/00295/2025), and FCS’s and JR’s CEEC individual contract (2024.09311.CEECIND and 2022.01763.CEECIND, respectively). HS acknowledges funding by the Deutsche Forschungsgemeinschaft (DFG, German Research Foundation) (grant 521529463).

