## Supporting Information for "Two centuries of change in the traits and origins of non-native vertebrates"

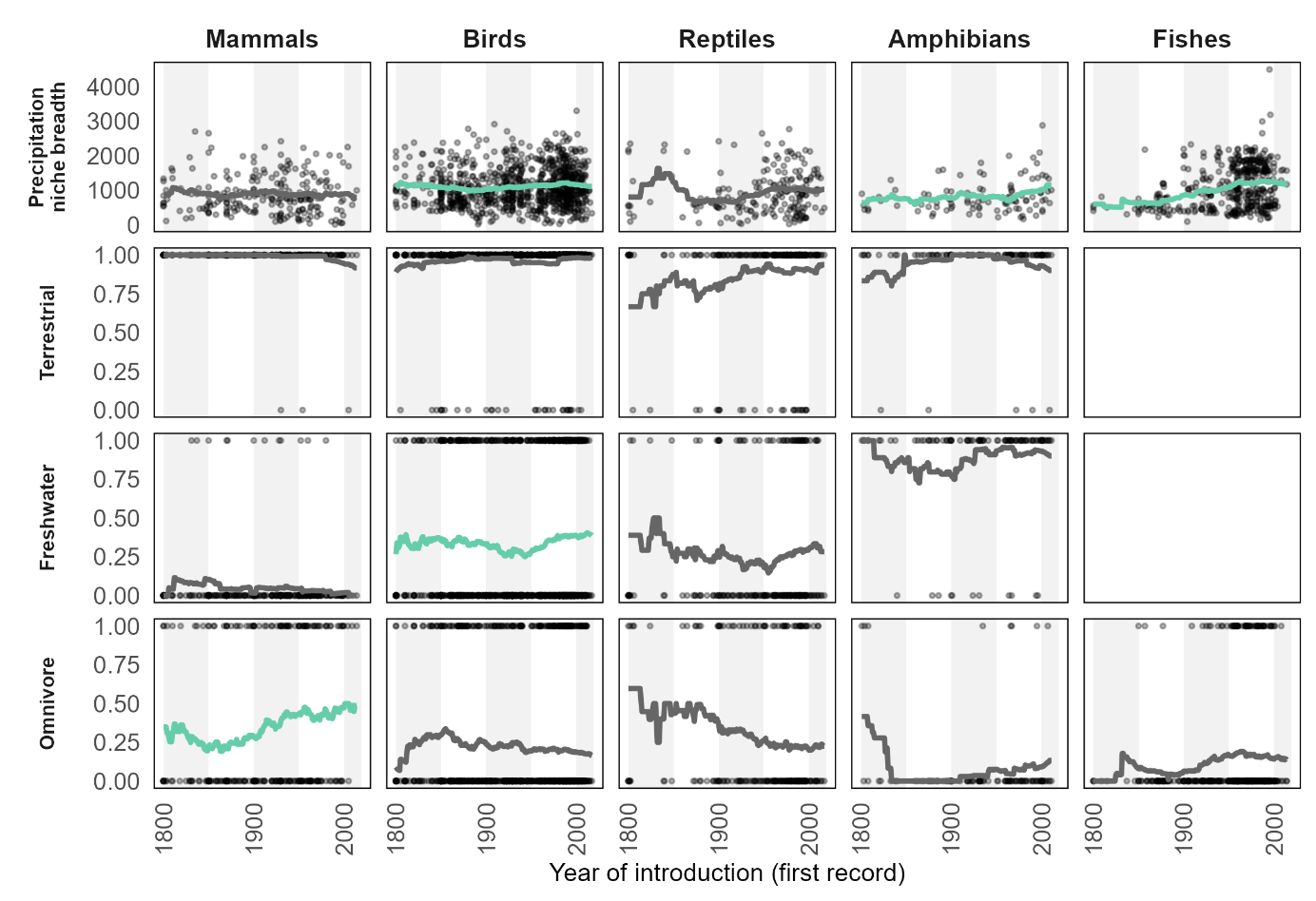
Figures

Fig. S1. Species-level trait values for the least consistent cross-taxon changes, plotted against the first-recorded year for each taxonomic group. Lines represent smoothed temporal trends based on a centered 50-year moving average of annual mean trait values, highlighting long-term patterns while reducing short-term variation. For binary traits, trends represent temporal changes in the proportion of newly recorded non-native species exhibiting the respective trait. Shaded vertical bands indicate 50-year periods used to visualize temporal structure. Points represent individual species according to their earliest known global first record. Line colors indicate the direction and significance of Kendall’s rank correlations: green denotes significant positive relationships, red denotes significant negative relationships, and black the non-significant relationships (statistical significance defined at *P* value <0.05). Some traits were log-transformed for visualization.

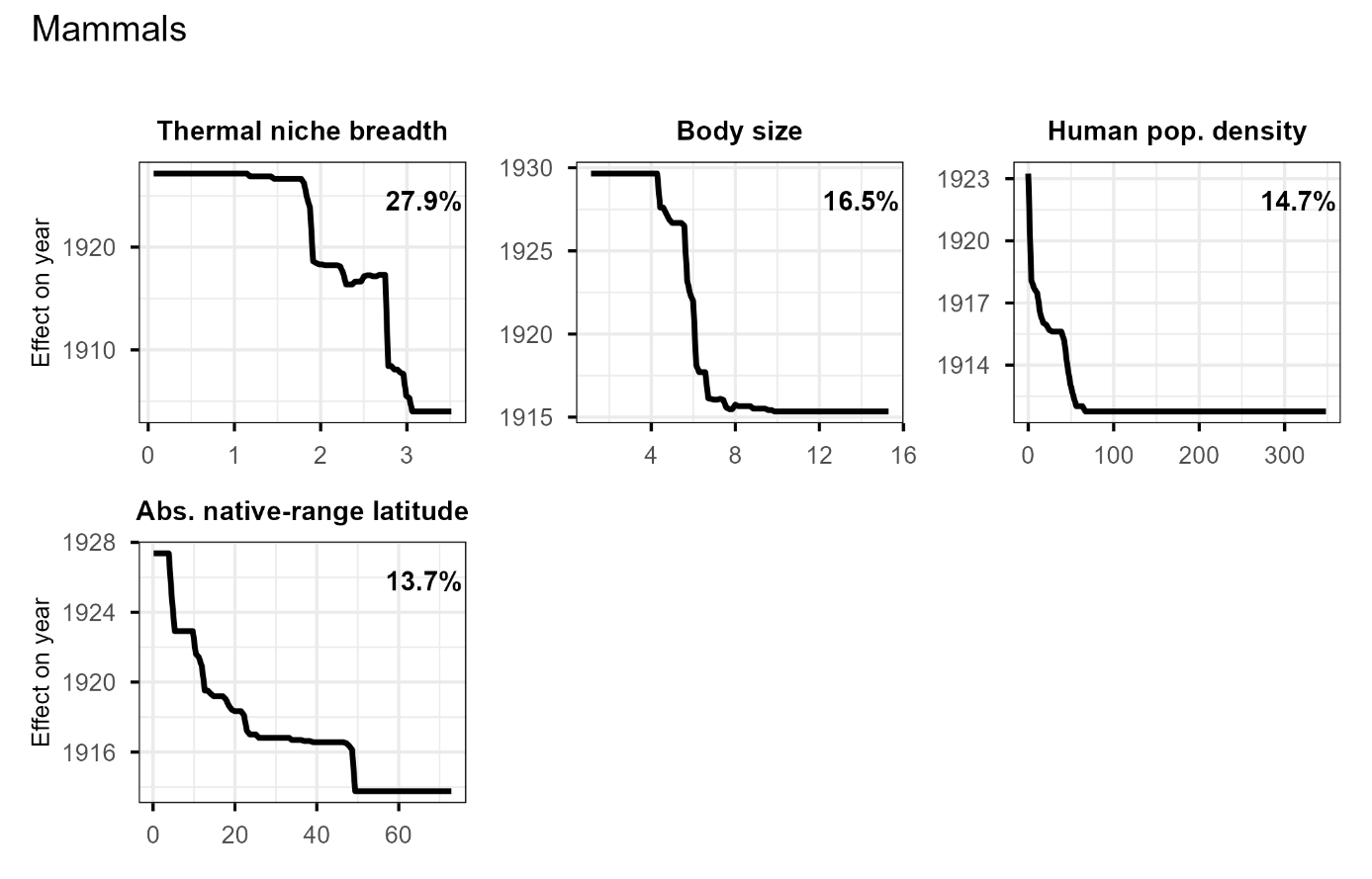

Fig. S2. Partial dependence plots for predictor variables with high relative influence (≥10%) derived from boosted regression tree models for mammals. Plots show the marginal effect of each trait on the timing of first recorded introductions (year), while averaging over the effects of all other predictors. Values on the y-axis represent the partial effect of each predictor on the modelled introduction year after accounting for regional random effects (i.e., the residual component of introduction year). Lower partial-effect values correspond to earlier modelled first-record years, whereas downward responses indicate traits associated with earlier introductions.

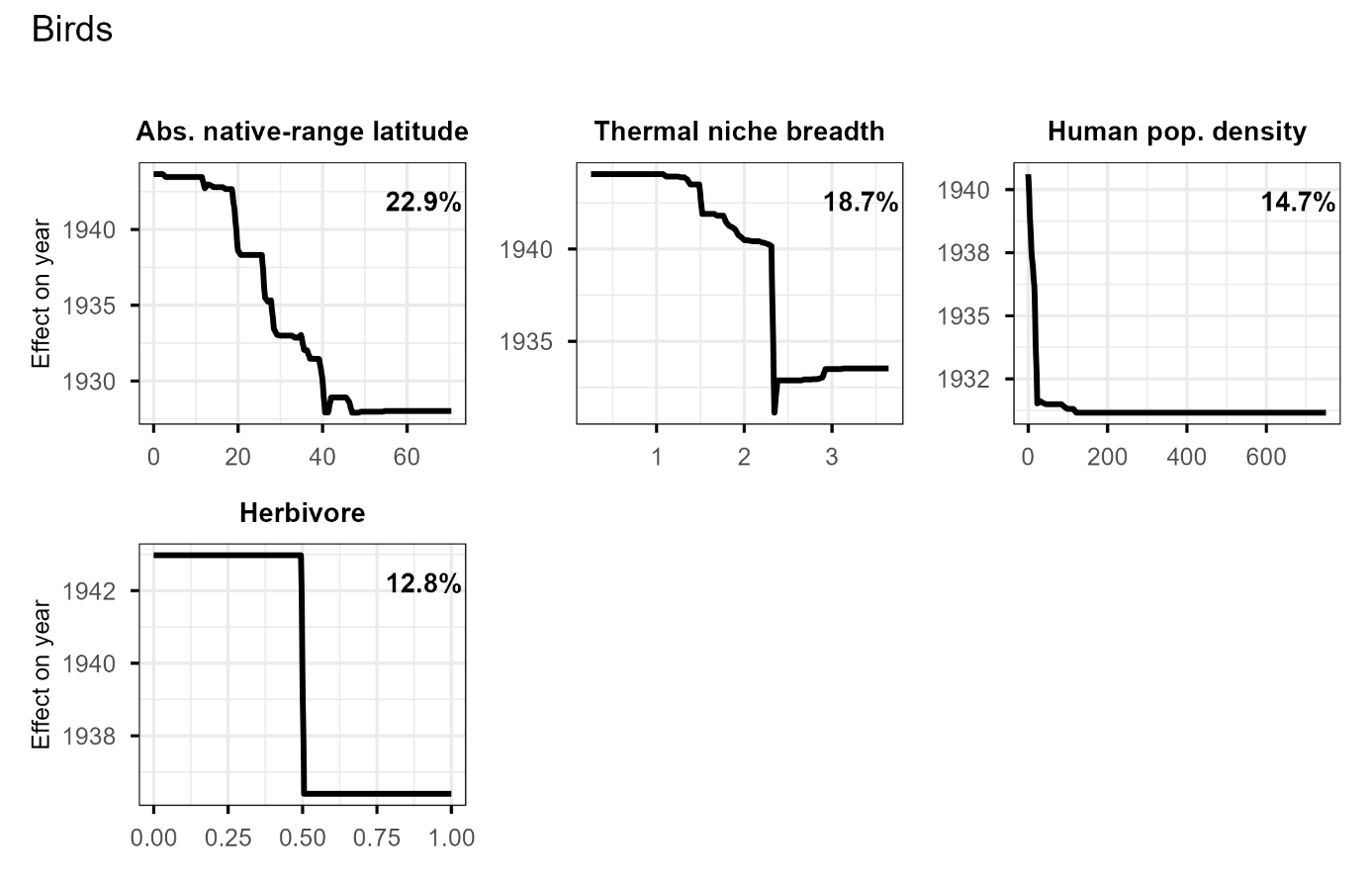
Fig. S3. Partial dependence plots for predictor variables with high relative influence (≥10%) derived from boosted regression tree models for birds. Plots show the marginal effect of each trait on the timing of first recorded introductions (year), while averaging over the effects of all other predictors. Values on the y-axis represent the partial effect of each predictor on the modelled introduction year after accounting for regional random effects (i.e., the residual component of introduction year). Lower partial-effect values correspond to earlier modelled first-record years, whereas downward responses indicate traits associated with earlier introductions.

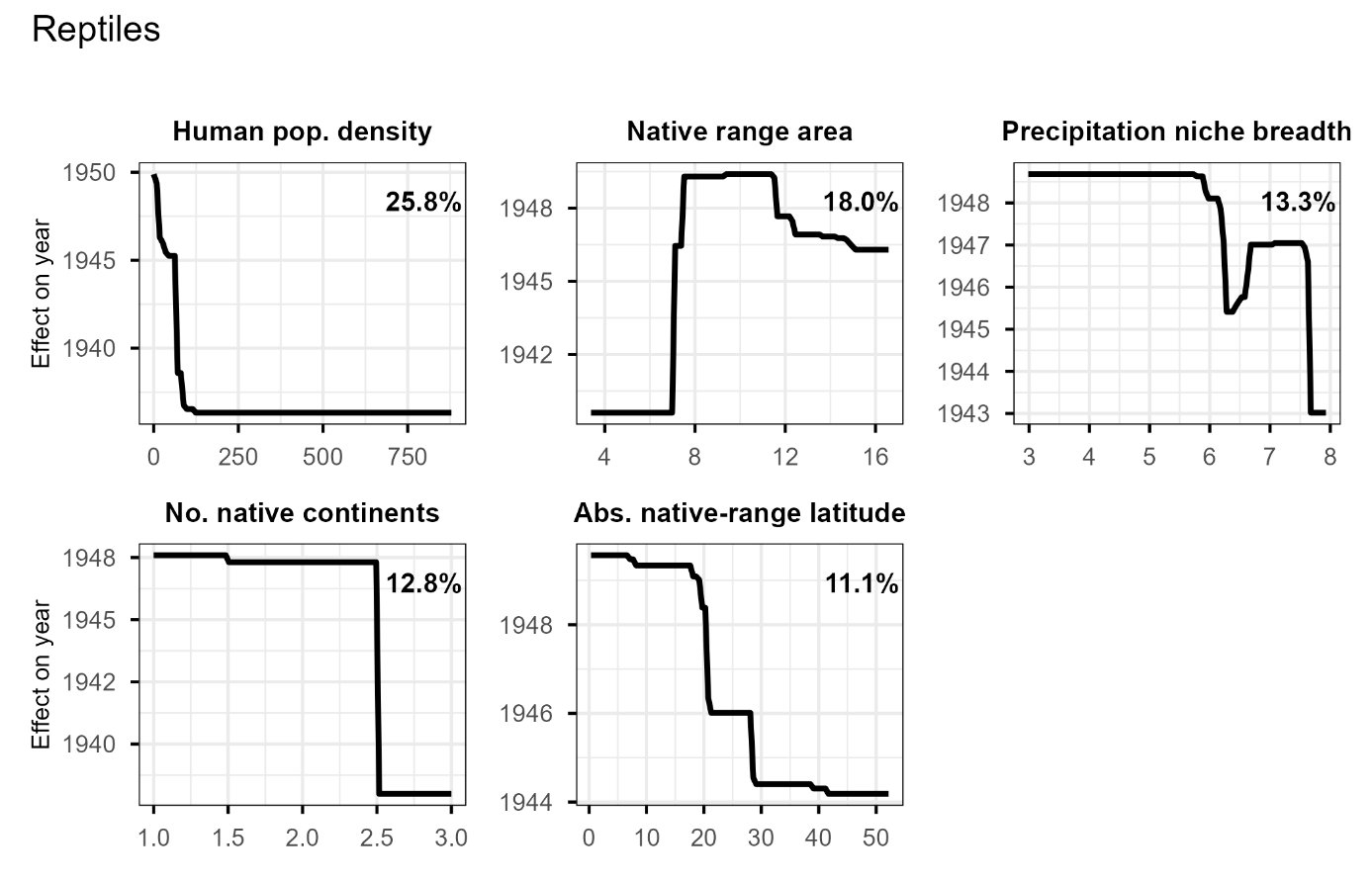
Fig. S4. Partial dependence plots for predictor variables with high relative influence (≥10%) derived from boosted regression tree models for reptiles. Plots show the marginal effect of each trait on the timing of first recorded introductions (year), while averaging over the effects of all other predictors. Values on the y-axis represent the partial effect of each predictor on the modelled introduction year after accounting for regional random effects (i.e., the residual component of introduction year). Lower partial-effect values correspond to earlier modelled first-record years, whereas downward responses indicate traits associated with earlier introductions.

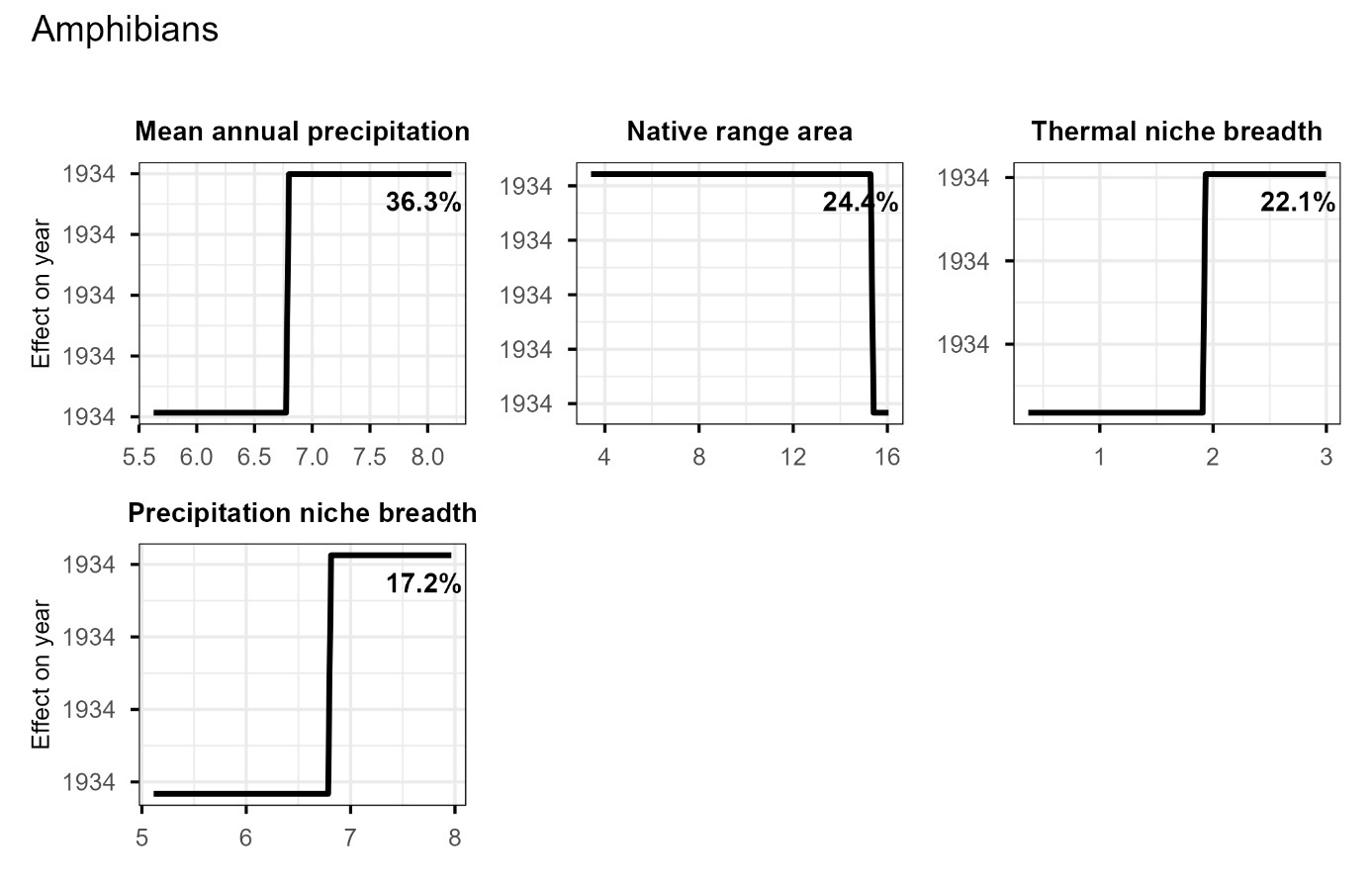
Fig. S5. Partial dependence plots for predictor variables with high relative influence (≥10%) derived from boosted regression tree models for amphibians. Plots show the marginal effect of each trait on the timing of first recorded introductions (year), while averaging over the effects of all other predictors. Values on the y-axis represent the partial effect of each predictor on the modelled introduction year after accounting for regional random effects (i.e., the residual component of introduction year). Lower partial-effect values correspond to earlier modelled first-record years, whereas downward responses indicate traits associated with earlier introductions.

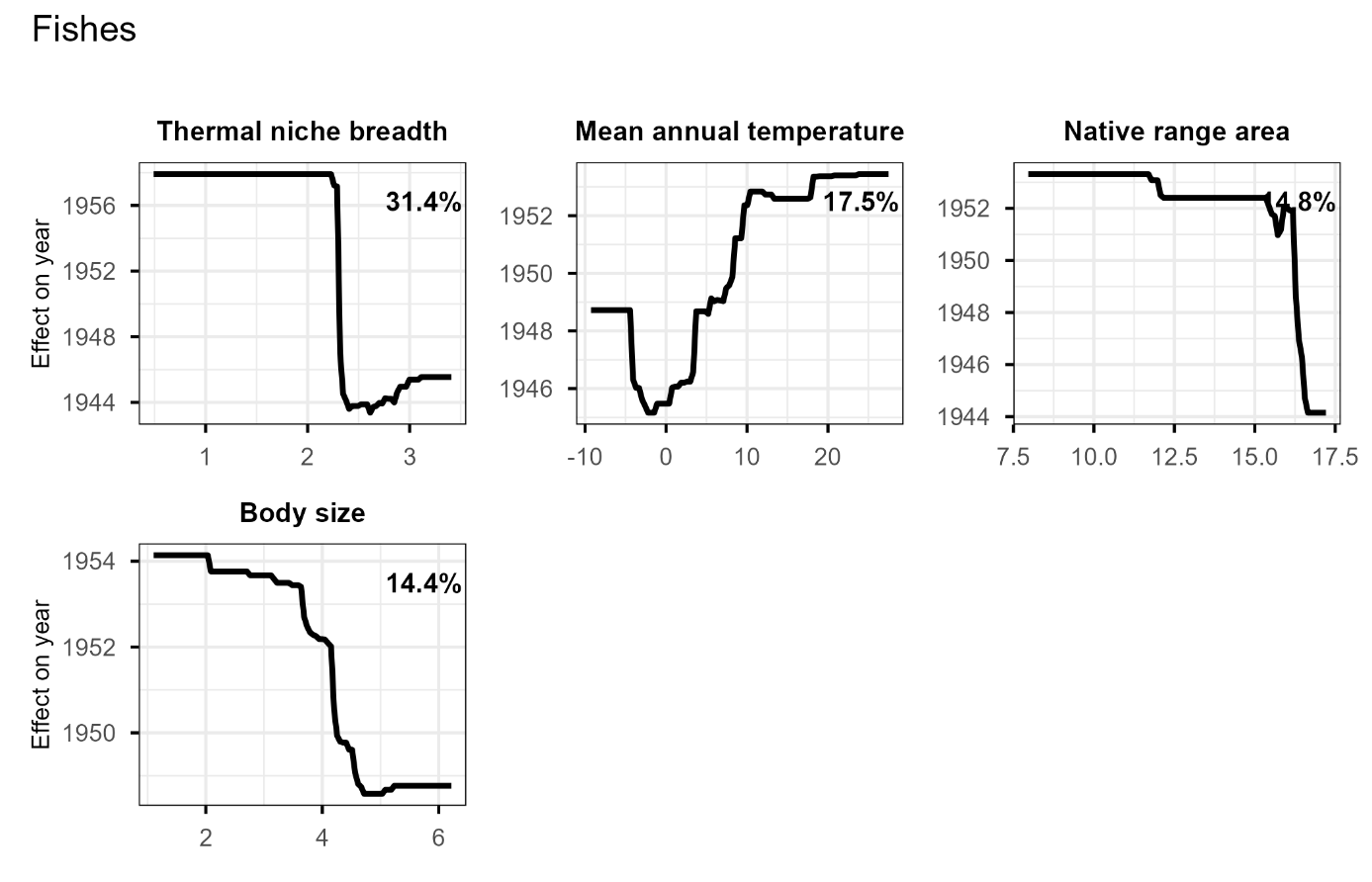
Fig. S6. Partial dependence plots for predictor variables with high relative influence (≥10%) derived from boosted regression tree models for fishes. Plots show the marginal effect of each trait on the timing of first recorded introductions (year), while averaging over the effects of all other predictors. Values on the y-axis represent the partial effect of each predictor on the modelled introduction year after accounting for regional random effects (i.e., the residual component of introduction year). Lower partial-effect values correspond to earlier modelled first-record years, whereas downward responses indicate traits associated with earlier introductions.

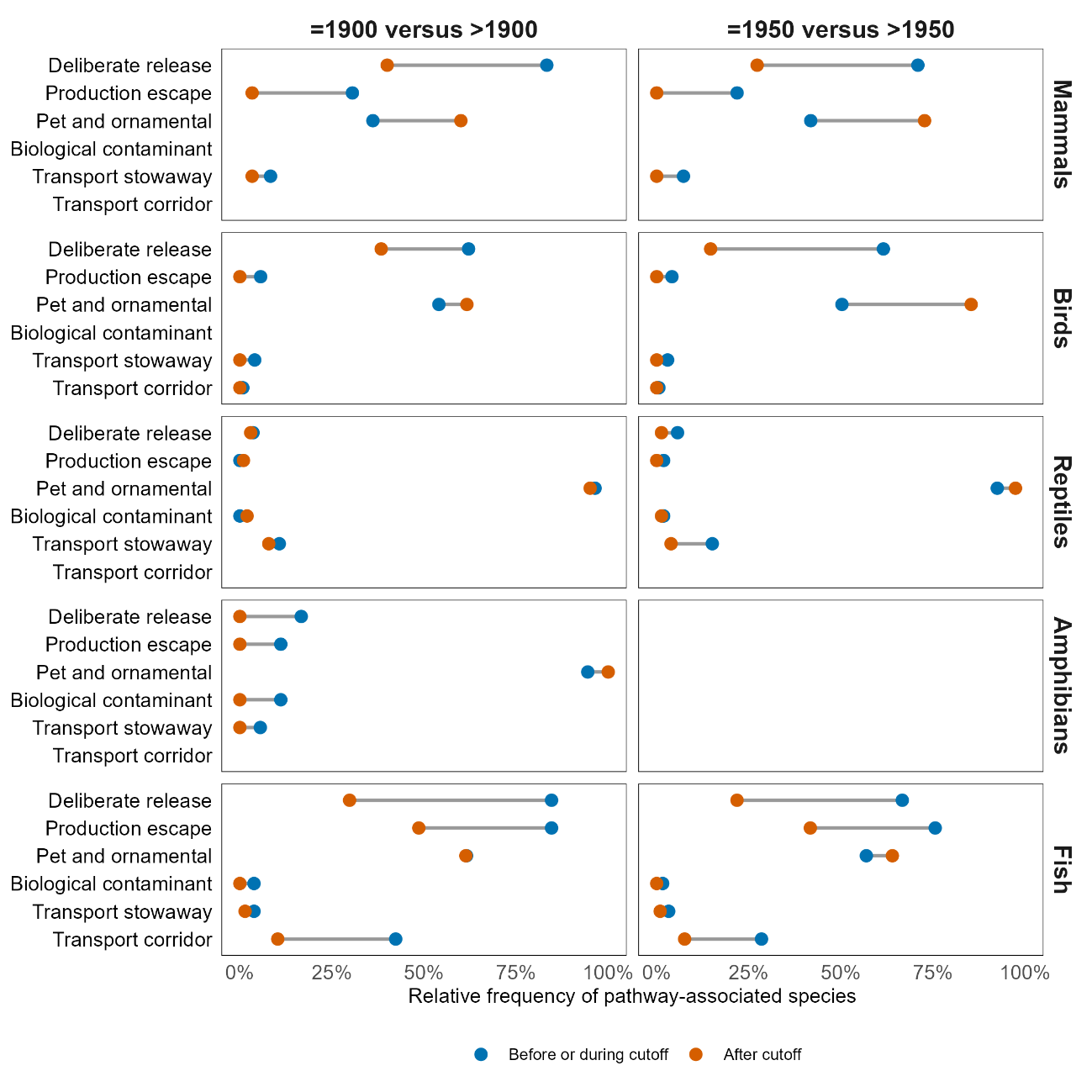
Fig. S7. Temporal changes in the relative frequency of introduction pathway classes across historical cut-offs. Relative frequency of species associated with each pathway class before/during, and after the 1900 and 1950 cut-offs. Blue points indicate species first recorded up to and including the cut-off, and orange points indicate species first recorded after the cut-off. Grey lines connect the two periods for each pathway, with wider lines indicating larger changes in pathway frequency.

Tables

Table S1. Functional and biogeographic traits used to characterize non-native species.

| **Variable name** | **Description** |
| --- | --- |
| Body size | Adult body size, measured as maximum body mass for amphibians and reptiles and mean body mass for birds and mammals, in grams; and as maximum body length for fishes, in centimeters. |
| Freshwater | Habitat use. Indicates whether the species primarily occupies freshwater habitats (binary: 0 = no, 1 = yes). |
| Terrestrial | Habitat use. Indicates whether the species primarily occupies terrestrial habitats (binary: 0 = no, 1 = yes). |
| Herbivore | Dietary guild. Indicates whether the species feeds mainly on plant material (binary: 0 = no, 1 = yes). |
| Carnivore | Dietary guild. Indicates whether the species feeds mainly on animal prey (binary: 0 = no, 1 = yes). |
| Omnivore | Dietary guild. Indicates whether the species feeds on both plant and animal material (binary: 0 = no, 1 = yes). |
| Native range area | Total area of the species' native geographic range (km²; continuous). |
| Absolute native range latitude | Absolute latitude of the geographic centroid of the native range (degrees; continuous). |
| Human population density | Median human population density within the native range in 1975 (people km⁻²; continuous). |
| Number of native continents | Total number of continents on which the species is native (continuous or integer). |
| Mean annual temperature | Median of mean annual temperature across the entire native range (°C; continuous). |
| Mean annual precipitation | Median of mean annual precipitation across the entire native range (mm; continuous). |
| Thermal niche breadth | Temperature niche breadth, calculated as the difference between the 90^th^ and 10^th^ percentiles of native range temperature values (°C; continuous). |
| Precipitation niche breadth | Precipitation niche breadth, calculated as the difference between the 90^th^ and 10^th^ percentiles of native range precipitation values (mm; continuous). |

Table S2. Kendall’s rank correlation coefficient (τ) and respective *P* values of each predictor for mammals (*N* = 234 pairs). Significant correlation coefficients are in bold (*P* value < 0.05).

| **Trait** | **τ coefficient** | ***P* values** |
| --- | --- | --- |
| **Body size** | **-0.104** | **0.018** |
| Freshwater | -0.050 | 0.351 |
| Terrestrial | -0.083 | 0.123 |
| Herbivore | 0.024 | 0.653 |
| **Carnivore** | **0.119** | **0.028** |
| **Omnivore** | **0.131** | **0.015** |
| **Native range area** | **-0.146** | **0.0009** |
| **Absolute native range latitude** | **-0.176** | **<0.0001** |
| Human population density | -0.039 | 0.382 |
| **Number of native continents** | **-0.181** | **0.0005** |
| **Mean annual temperature** | **0.144** | **0.001** |
| Mean annual precipitation | 0.050 | 0.257 |
| **Thermal niche breadth** | **-0.216** | **<0.0001** |
| Precipitation niche breadth | -0.017 | 0.698 |

Table S3. Kendall’s rank correlation coefficient (τ) and respective *P* values of each predictor for birds (*N* = 931 pairs). Significant correlation coefficients are in bold (*P* value < 0.05).

| **Trait** | **τ coefficient** | ***P* values** |
| --- | --- | --- |
| Body size | -0.005 | 0.817 |
| **Freshwater** | **0.076** | **0.005** |
| Terrestrial | 0.020 | 0.449 |
| **Herbivore** | **-0.153** | **<0.001** |
| **Carnivore** | **0.107** | **<0.001** |
| Omnivore | -0.027 | 0.317 |
| **Native range area** | **-0.057** | **0.009** |
| **Absolute native range latitude** | **-0.151** | **<0.001** |
| Human population density | 0.018 | 0.412 |
| **Number of native continents** | **-0.103** | **<0.001** |
| **Mean annual temperature** | **0.118** | **<0.001** |
| **Mean annual precipitation** | **0.078** | **<0.001** |
| **Thermal niche breadth** | **-0.088** | **<0.001** |
| **Precipitation niche breadth** | **0.054** | **0.015** |

Table S4. Kendall’s rank correlation coefficient (τ) and respective *P* values of each predictor for reptiles (*N* = 198 pairs). Significant correlation coefficients are in bold (*P* value < 0.05).

| **Trait** | **τ coefficient** | ***P* values** |
| --- | --- | --- |
| Body size | 0.043 | 0.371 |
| Freshwater | 0.060 | 0.305 |
| Terrestrial | 0.034 | 0.560 |
| **Herbivore** | **-0.116** | **0.047** |
| Carnivore | 0.029 | 0.618 |
| Omnivore | -0.106 | 0.072 |
| Native range area | 0.005 | 0.915 |
| **Absolute native range latitude** | **-0.131** | **0.006** |
| **Human population density** | **-0.110** | **0.021** |
| Number of native continents | -0.055 | 0.338 |
| **Mean annual temperature** | **0.111** | **0.021** |
| Mean annual precipitation | 0.072 | 0.136 |
| Thermal niche breadth | -0.025 | 0.604 |
| Precipitation niche breadth | 0.073 | 0.128 |

Table S5. Kendall’s rank correlation coefficient (τ) and respective *P* values of each predictor for amphibians (*N* = 100 pairs). Significant correlation coefficients are in bold (*P* value < 0.05).

| **Trait** | **τ coefficient** | ***P* values** |
| --- | --- | --- |
| Body size | -0.092 | 0.176 |
| Freshwater | 0.062 | 0.454 |
| Terrestrial | -0.024 | 0.776 |
| Herbivore | -0.040 | 0.629 |
| Omnivore | -0.040 | 0.629 |
| Native range area | -0.053 | 0.437 |
| Absolute native range latitude | -0.117 | 0.086 |
| Human population density | 0.078 | 0.255 |
| Number of native continents | -0.008 | 0.923 |
| Mean annual temperature | 0.083 | 0.224 |
| **Mean annual precipitation** | **0.150** | **0.028** |
| Thermal niche breadth | 0.045 | 0.510 |
| **Precipitation niche breadth** | **0.140** | **0.040** |

Table S6. Kendall’s rank correlation coefficient (τ) and respective *P* values of each predictor for fishes (*N* = 445 pairs). Significant correlation coefficients are in bold (*P* value < 0.05).

| **Trait** | **τ coefficient** | ***P* values** |
| --- | --- | --- |
| **Body size** | **-0.168** | **<0.0001** |
| **Herbivore** | **0.103** | **0.008** |
| **Carnivore** | **-0.090** | **0.021** |
| Omnivore | 0.055 | 0.156 |
| **Native range area** | **-0.144** | **<0.0001** |
| **Absolute native range latitude** | **-0.312** | **<0.0001** |
| Human population density | 0.020 | 0.543 |
| **Number of native continents** | **-0.244** | **<0.0001** |
| **Mean annual temperature** | **0.283** | **<0.0001** |
| **Mean annual precipitation** | **0.276** | **<0.0001** |
| **Thermal niche breadth** | **-0.227** | **<0.0001** |
| **Precipitation niche breadth** | **0.162** | **<0.0001** |

Table S7. Variance inflation factors (VIF) for each predictor across taxonomic groups. Predictors exhibiting VIF values greater than 10 were removed.

| **Trait** | **Mammals** | **Birds** | **Reptiles** | **Amphibians** | **Fishes** |
| --- | --- | --- | --- | --- | --- |
| Body size | 1.601 | 1.127 | 1.851 | 1.335 | 1.189 |
| Freshwater | 1.109 | 1.197 | 2.091 | 1.222 | - |
| Terrestrial | 1.157 | 1.138 | 1.867 | 1.094 | - |
| Herbivore | 1.441 | 1.190 | 1.375 | 1.202 | - |
| Carnivore | - | - | 1.606 | - | 1.055 |
| Omnivore | 1.571 | 1.144 | - | - | 1.075 |
| Native range area | 2.586 | 2.649 | 3.877 | 2.285 | 1.918 |
| Absolute native range latitude | 8.991 | 6.503 | 6.101 | - | - |
| Human population density | 1.257 | 1.133 | 1.289 | 1.341 | 1.259 |
| Number of native continents | 1.353 | 1.374 | 1.235 | 1.214 | 1.286 |
| Mean annual temperature | 8.990 | 7.789 | 6.285 | 2.096 | 4.513 |
| Mean annual precipitation | 3.180 | 2.960 | 2.958 | 1.871 | 3.481 |
| Thermal niche breadth | 3.242 | 3.927 | 3.118 | 3.049 | 2.035 |
| Precipitation niche breadth | 5.087 | 2.699 | 5.172 | 1.983 | 4.012 |

Table S8. Results of LMM evaluating the relationship between species traits and the year of first recorded introduction for mammals. Fixed effects include functional and biogeographic traits, with estimated (coefficients), standard errors and associated *P* values. Variance components and standard deviations (SD) are reported for random effects (including continent). Marginal R^2^ represents the variance explained by fixed effects, and conditional R^2^ the proportion of variance explained by both fixed and random effects. Significant coefficients are in bold (*P* value < 0.05).

| **Trait** | **Coefficient** | **Standard error** | ***P* values** |
| --- | --- | --- | --- |
| Body size | -5.573 | 3.895 | 0.154 |
| Freshwater | 0.138 | 3.190 | 0.966 |
| Terrestrial | -5.253 | 3.255 | 0.108 |
| Herbivore | -5.960 | 3.674 | 0.106 |
| **Omnivore** | **8.938** | **3.872** | **0.022** |
| Native range area | -1.930 | 4.959 | 0.698 |
| **Absolute native range latitude** | **-35.181** | **9.240** | **0.0002** |
| Human population density | -3.913 | 3.437 | 0.256 |
| **Number of native continents** | **-12.283** | **3.574** | **0.0007** |
| **Mean annual temperature** | **-26.955** | **9.268** | **0.004** |
| Mean annual precipitation | 0.011 | 5.442 | 0.998 |
| **Thermal niche breadth** | **-11.859** | **5.457** | **0.031** |
| Precipitation niche breadth | -8.893 | 6.870 | 0.197 |
| Random effects (variance, SD) | Continent: 122, 11.05  Residual: 2076, 45.56 | | |
| R^2^ marginal: 0.276; R^2^ conditional: 0.316 | | | |

Table S9. Results of LMM evaluating the relationship between species traits and the year of first recorded introduction for birds. Fixed effects include functional and biogeographic traits, with estimated (coefficients), standard errors and associated *P* values. Variance components and standard deviations (SD) are reported for random effects (including continent). Marginal R^2^ represents the variance explained by fixed effects, and conditional R^2^ the proportion of variance explained by both fixed and random effects. Carnivore trait was removed due to a high VIF value (VIF = 234.04). Significant coefficients are in bold (*P* value < 0.05).

| **Trait** | **Coefficient** | **Standard error** | ***P* values** |
| --- | --- | --- | --- |
| **Body size** | **-4.860** | **1.696** | **0.004** |
| **Freshwater** | **4.632** | **1.688** | **0.006** |
| Terrestrial | 0.466 | 1.625 | 0.774416 |
| **Herbivore** | **-10.440** | **1.675** | **<0.0001** |
| Omnivore | 2.892 | 1.637 | 0.078 |
| **Native range area** | **-6.027** | **2.497** | **0.016** |
| **Absolute native range latitude** | **-21.943** | **3.975** | **<0.0001** |
| **Human population density** | **-3.746** | **1.633** | **0.022** |
| **Number of native continents** | **-6.878** | **1.793** | **0.0001** |
| **Mean annual temperature** | **-13.229** | **4.301** | **0.002** |
| **Mean annual precipitation** | **-6.656** | **2.649** | **0.012** |
| **Thermal niche breadth** | **-8.278** | **3.038** | **0.007** |
| Precipitation niche breadth | 2.049 | 2.505 | 0.414 |
| Random effects (variance, SD) | Continent: 755.3, 27.48  Residual: 2132.6, 46.18 | | |
| R^2^ marginal: 0.130; R^2^ conditional: 0.358 | | | |

Table S10. Results of LMM evaluating the relationship between species traits and the year of first recorded introduction for reptiles. Fixed effects include functional and biogeographic traits, with estimated (coefficients), standard errors and associated *P* values. Variance components and standard deviations (SD) are reported for random effects (including continent). Marginal R^2^ represents the variance explained by fixed effects, and conditional R^2^ the proportion of variance explained by both fixed and random effects. Significant coefficients are in bold (*P* value < 0.05).

| **Trait** | **Coefficient** | **Standard error** | ***P* values** |
| --- | --- | --- | --- |
| Body size | -1.634 | 4.678 | 0.727 |
| Freshwater | 3.014 | 4.950 | 0.543 |
| Terrestrial | 4.415 | 4.673 | 0.346 |
| Herbivore | -4.869 | 4.045 | 0.230 |
| Carnivore | -2.020 | 4.321 | 0.641 |
| Native range area | 1.441 | 6.721 | 0.830 |
| **Absolute native range latitude** | **-30.423** | **8.505** | **0.0004** |
| Human population density | -1.280 | 3.914 | 0.744 |
| **Number of native continents** | **-12.162** | **3.975** | **0.003** |
| Mean annual temperature | -14.299 | 8.773 | 0.105 |
| **Mean annual precipitation** | **-12.080** | **6.052** | **0.047** |
| Thermal niche breadth | -0.890 | 6.104 | 0.884 |
| Precipitation niche breadth | 4.348 | 7.853 | 0.581 |
| Random effects (variance, SD) | Continent: 537.3, 23.18  Residual: 2222.8, 47.15 | | |
| R^2^ marginal: 0.131; R^2^ conditional: 0.300 | | | |

Table S11. Results of LMM evaluating the relationship between species traits and the year of first recorded introduction for amphibians. Fixed effects include functional and biogeographic traits, with estimated (coefficients), standard errors and associated *P* values. Variance components and standard deviations (SD) are reported for random effects (including continent). Marginal R^2^ represents the variance explained by fixed effects, and conditional R^2^ the proportion of variance explained by both fixed and random effects. Significant coefficients are in bold (*P* value < 0.05).

| **Trait** | **Coefficient** | **Standard error** | ***P* values** |
| --- | --- | --- | --- |
| Body size | -6.780 | 6.422 | 0.294 |
| Freshwater | -0.698 | 6.030 | 0.908 |
| Terrestrial | 1.027 | 5.653 | 0.856 |
| Herbivore | -7.258 | 5.959 | 0.227 |
| Native range area | -13.611 | 8.244 | 0.102 |
| Human population density | 9.001 | 6.331 | 0.159 |
| Number of native continents | 9.071 | 6.404 | 0.161 |
| Mean annual temperature | 9.154 | 8.176 | 0.266 |
| Mean annual precipitation | 9.439 | 7.540 | 0.214 |
| **Thermal niche breadth** | **30.429** | **9.460** | **0.002** |
| Precipitation niche breadth | -1.641 | 7.629 | 0.830 |
| Random effects (variance, SD) | Continent: 237.3, 15.40  Residual: 2826.7, 53.17 | | |
| R^2^ marginal: 0.165; R^2^ conditional: 0.230 | | | |

Table S12. Results of LMM evaluating the relationship between species traits and the year of first recorded introduction for fishes. Fixed effects include functional and biogeographic traits, with estimated (coefficients), standard errors and associated *P* values. Variance components and standard deviations (SD) are reported for random effects (including continent). Marginal R^2^ represents the variance explained by fixed effects, and conditional R^2^ the proportion of variance explained by both fixed and random effects. Significant coefficients are in bold (*P* value < 0.05).

| **Trait** | **Coefficient** | **Standard error** | ***P* values** |
| --- | --- | --- | --- |
| **Body size** | **-4.374** | **1.802** | **0.016** |
| Carnivore | -1.900 | 1.682 | 0.259 |
| Omnivore | 1.399 | 1.710 | 0.414 |
| Native range area | -2.544 | 2.275 | 0.264 |
| Human population density | -0.984 | 1.844 | 0.594 |
| **Number of native continents** | **-6.769** | **1.874** | **0.0003** |
| **Mean annual temperature** | **14.558** | **3.662** | **<0.0001** |
| Mean annual precipitation | -2.557 | 3.101 | 0.410 |
| Thermal niche breadth | -3.268 | 2.356 | 0.166 |
| Precipitation niche breadth | -0.522 | 3.292 | 0.874 |
| Random effects (variance, SD) | Continent: 89.46, 9.458  Residual: 1183.74, 34.405 | | |
| R^2^ marginal: 0.257; R^2^ conditional: 0.309 | | | |

Table S13. Relative influence (%) of predictor variables obtained from boosted regression tree models for mammals. Values represent the contribution of each trait to explaining variation in the timing of first recorded introductions. Model predictive performance is reported as relative absolute error (RAE = 0.895).

| **Trait** | **Variable importance (%)** |
| --- | --- |
| Body size | 16.527 |
| Freshwater | 0 |
| Terrestrial | 0 |
| Herbivore | 0 |
| Omnivore | 2.792 |
| Native range area | 7.414 |
| Absolute native range latitude | 13.685 |
| Human population density | 14.741 |
| Number of native continents | 7.956 |
| Mean annual temperature | 3.245 |
| Mean annual precipitation | 3.494 |
| Thermal niche breadth | 27.950 |
| Precipitation niche breadth | 1.438 |

Table S14. Relative influence (%) of predictor variables obtained from boosted regression tree models for birds. Values represent the contribution of each trait to explaining variation in the timing of first recorded introductions. Model predictive performance is reported as relative absolute error (RAE = 0.798).

| **Trait** | **Variable importance (%)** |
| --- | --- |
| Body size | 5.279 |
| Freshwater | 0.673 |
| Terrestrial | 0 |
| Herbivore | 12.842 |
| Omnivore | 0 |
| Native range area | 8.648 |
| Absolute native range latitude | 22.851 |
| Human population density | 14.740 |
| Number of native continents | 9.369 |
| Mean annual temperature | 1.561 |
| Mean annual precipitation | 1.741 |
| Thermal niche breadth | 18.656 |
| Precipitation niche breadth | 1.883 |

Table S15. Relative influence (%) of predictor variables obtained from boosted regression tree models for reptiles. Values represent the contribution of each trait to explaining variation in the timing of first recorded introductions. Model predictive performance is reported as relative absolute error (RAE = 0.91).

| **Trait** | **Variable importance (%)** |
| --- | --- |
| Body size | 3.741 |
| Freshwater | 0.114 |
| Terrestrial | 0 |
| Herbivore | 1.505 |
| Omnivore | 0.703 |
| Native range area | 17.972 |
| Absolute native range latitude | 11.100 |
| Human population density | 25.787 |
| Number of native continents | 12.796 |
| Mean annual temperature | 5.818 |
| Mean annual precipitation | 2.273 |
| Thermal niche breadth | 4.907 |
| Precipitation niche breadth | 13.283 |

Table S16. Relative influence (%) of predictor variables obtained from boosted regression tree models for amphibians. Values represent the contribution of each trait to explaining variation in the timing of first recorded introductions. Model predictive performance is reported as relative absolute error (RAE = 0.97).

| **Trait** | **Variable importance (%)** |
| --- | --- |
| Body size | 0 |
| Freshwater | 0 |
| Terrestrial | 0 |
| Herbivore | 0 |
| Native range area | 24.368 |
| Human population density | 0 |
| Number of native continents | 0 |
| Mean annual temperature | 0 |
| Mean annual precipitation | 36.377 |
| Thermal niche breadth | 22.141 |
| Precipitation niche breadth | 17.164 |

Table S17. Relative influence (%) of predictor variables obtained from boosted regression tree models for fishes. Values represent the contribution of each trait to explaining variation in the timing of first recorded introductions. Model predictive performance is reported as relative absolute error (RAE = 0.827).

| **Trait** | **Variable importance (%)** |
| --- | --- |
| Body size | 14.436 |
| Carnivore | 0 |
| Omnivore | 0 |
| Native range area | 14.811 |
| Human population density | 2.610 |
| Number of native continents | 1.906 |
| Mean annual temperature | 17.453 |
| Mean annual precipitation | 8.938 |
| Thermal niche breadth | 31.385 |
| Precipitation niche breadth | 8.460 |
